# Pathogen dependence and inter-individual variability of post-infection reproductive fitness in *Drosophila melanogaster*

**DOI:** 10.1101/2022.05.22.492957

**Authors:** Aabeer Basu, Vandana Gupta, Nagaraj Guru Prasad

## Abstract

In the experiments reported in this manuscript, we explore the effect of bacterial infections on the reproductive output of *Drosophila melanogaster* females. Canonical view of host-pathogen interactions supposes two possible outcomes. Because of immune defence being an energy/resource intensive function, an infected female reallocates resources away from reproductive processes and towards immune defence, therefore compromising its reproductive output. Alternatively, faced with impending mortality, an infected female increases its reproductive output to compensate for lost opportunities of future reproduction. We tested if pathogen identity, infection outcome (survival vs. death), and/or time of death determines the reproductive output of females infected with three bacterial pathogens. Our results show that pathogen identity is a reliable predictor of population level response of infected females but does not reliably predict the behaviour of individual females. Additionally, females succumbing to infection exhibit greater variability in reproductive output, compared to both survivors and controls, but this variability is not explained by either the time of death or the identity of the infecting pathogen. Furthermore, survivors of infection have reproductive output similar to control females.

## INTRODUCTION

Omnipresence of pathogens/parasites impose a strong selection pressure on hosts to evolve mechanisms of defense. Such defense mechanisms go far beyond the canonical anatomical and physiological defenses, and include behavioral strategies that either help alleviate risk of infection or help mitigate the consequences. Fecundity compensation, that is the post infection increase in reproductive effort of the host, is one such behavioral defense that helps hosts maintain their evolutionary fitness [1]. Since increased reproductive effort maximizes immediate reproductive output at the cost of future chance of reproduction [2], organisms under benign conditions are expected to pace out their reproductive schedule so as to maximize their life-time reproductive success [3, 4]. Under circumstances which lead to pre-mature death, such as a lethal infection, future opportunities of reproduction are compromised and organisms should, in theory, maximize their immediate reproductive effort [5, 6, 7]. Minchella and Loverde [8] first demonstrated this phenomenon in snails infected with castrating trematode parasites, where hosts increased their immediate reproductive output in response to parasitic infection.

An infection is also detrimental to the physiology of the host organism. One, mounting an immune response requires investing energy and resources that could otherwise have been utilized elsewhere, such as towards reproduction [9, 10, 11]. Two, infection leads to somatic damage caused by the virulence factors produced by parasites/pathogens [12, 13]. And three, the immune response mounted by the host often causes collateral somatic damage to the host, leading to immunopathology [14]. Altogether this suggests that post-infection fitness of hosts depends on its ability to restrict the systemic propagation of the parasite/pathogen, plus the host’s capacity to continue to maintain physiological functionality during and after recovery from the infection [15]. Reallocating resources towards mounting an immune defense can lead to reduced reproductive effort during acute infection [16], and lingering somatic damage can keep reproductive effort to a minimum even after recovery. Fecundity compensation, as described above, therefore might not be the observed strategy in case of all hosts on every occasion, and will depend on the features of the specific host-pathogen system being studied.

The choice of strategy is likely to depend on the balance between the actual risk of mortality and the level of somatic damage incurred by the host. A greater risk of mortality should induce a stronger fecundity compensation response, thereby increasing reproductive effort, while reproductive effort should decline proportionately with increasing somatic damage. This balance can vary at the level of individual hosts, causing the mean population behavior to not be a true reflection of the individual variation in strategies. In fact, increased and decreased reproductive effort can be viewed as two ends of a continuum – instead of a dichotomous choice – with each individual host opting for an optimal level of reproductive effort based on their proximate circumstances.

Furthermore, post-infection reduction in host reproductive effort may also be driven by leeching of resources by the pathogen/parasite, damage to reproductive tissue, or manipulation of the host physiology by virulence factors produced by the pathogen/parasite [12]. Thus, post-infection reduction in host reproductive effort can also be a consequence of the infection process (presence of pathogen), independent of the host response to infection. To differentiate between post-infection phenotypes that are driven by pathogen manipulation and those caused by host immune response, previous studies have often used attenuated pathogens or pathogen-like proxies (bacteria-derived lipopolysaccharides, plastic beads, etc.; viz. [17]), arguing that such proxies stimulate the host immune system without causing any infection-related pathologies. While experiments with live pathogens may fail to tease apart host’s response to pathogens from pathogen’s manipulation of the host, experiments with pathogen-like proxies may not induce any fitness effects, both physiological and reproductive. Furthermore, given that mounting an immune response is costly to the host, hosts are under pressure to evolve mechanisms that differentiate real infections from false alerts. Thus, results obtained from experiments using attenuated pathogens and proxies are difficult to interpret. The results of experiments are thus likely to depend upon the exact biology of the interacting host and pathogen [18], on the physiological capability of the host to modify its own reproductive effort, and on whether such modifications of reproductive effort will materialize into benefits in terms of immune function [19], among various other factors.

Previous studies exploring the effect of parasites and pathogens on reproductive behavior of *Drosophila melanogaster* have reported diverse outcomes, depending partially upon the type of infectious agent used in the experiments. Flies having successfully survived a parasitoid attack as larvae have reduced fecundity as adults [20, 21]. Flies infected with Drosophila C Virus exhibit genotype and infection route dependent increase or decrease in reproductive output [22]. Infection with bacterial pathogens have been demonstrated to increase [23], reduce [24, 25], or maintain fecundity at an unaltered level [26, 27]. The reasons for this diversity of outcomes can be multiple, including host susceptibility to pathogens used [28], infection route [29, 30], genotypic differences in host strains and possible interactions with environmental factors [31, 32]. Another variable that can affect experimental outcomes is whether reproductive effort is measured during the acute or the chronic phase of infection [16]. Infection survivors continue to have a low level of systemic pathogen presence which have life history consequences [33], although one study reported infection survivors have similar fecundity as to the controls following recovery from a bacterial infection [34].

In this study we challenged *Drosophila melanogaster* females with three pathogenic bacteria, (a) *Bacillus thuringiensis*, (b) *Pseudomonas aeruginosa*, and (c) *Seratia marscesens*, and quantified their change in post-infection reproductive output, during the acute phase of infection, compared to uninfected controls. The aim of the study was to identify the effect of (a) pathogen identity, (b) infection outcome, and (c) time of death, on post-infection reproductive effort. Pathogen identity represents differences in pathogen virulence factors, host defence mechanisms, associated costs and immunopathology. Therefore, we expect that pathogen identity will be a strong determining factor for post-infection reproductive effort. Infection outcome, that is survival versus death, is the ultimate determinant of fitness at the level of individual hosts. Hosts that succumb to infection lose out on future opportunities to reproduce, and therefore are expected to modulate their current reproductive effort differently than hosts that recover from infection. And finally, individual hosts that die within a short period following infection are expected to exhibit a greater increase in reproductive effort compared to hosts that die relatively later.

## RESULTS

Using flies from a wild-type, outbred population of *Drosophila melanogaster* (BRB2, see MATERIALS AND METHODS for more details) we tested for the effect of infection with different entomopathogenic bacteria on female reproductive fitness. In the first experiment (figure 1.A), we infected 4-5-day old, inseminated females with three bacteria: *Bacillus thuringiensis* (hereafter *Bt*), *Pseudomonas aeruginosa* (hereafter *Pa*), and *Serratia marcescens* (hereafter *Sm*); we maintained sham-infected and uninfected controls along with the infected treatments. After infections, the females of each treatment were hosted in vials in groups of 8, with 10 vials per treatment. The entire experiment was independently replicated thrice. We monitored the mortality in these vials, every 2 hours, for 24 hours post-infection, covering the acute phase of infection of all three pathogens. As a measure of reproductive output, we counted the total number of eggs in each vial, laid by 8 females in 24 hours, and also the total number of adult progeny that developed from the eggs. This provided us with an additional measure of fitness: pre-adult viability (proportion of eggs that successfully developed into adults) of the progeny produced by the infected females.

**Figure 1.**
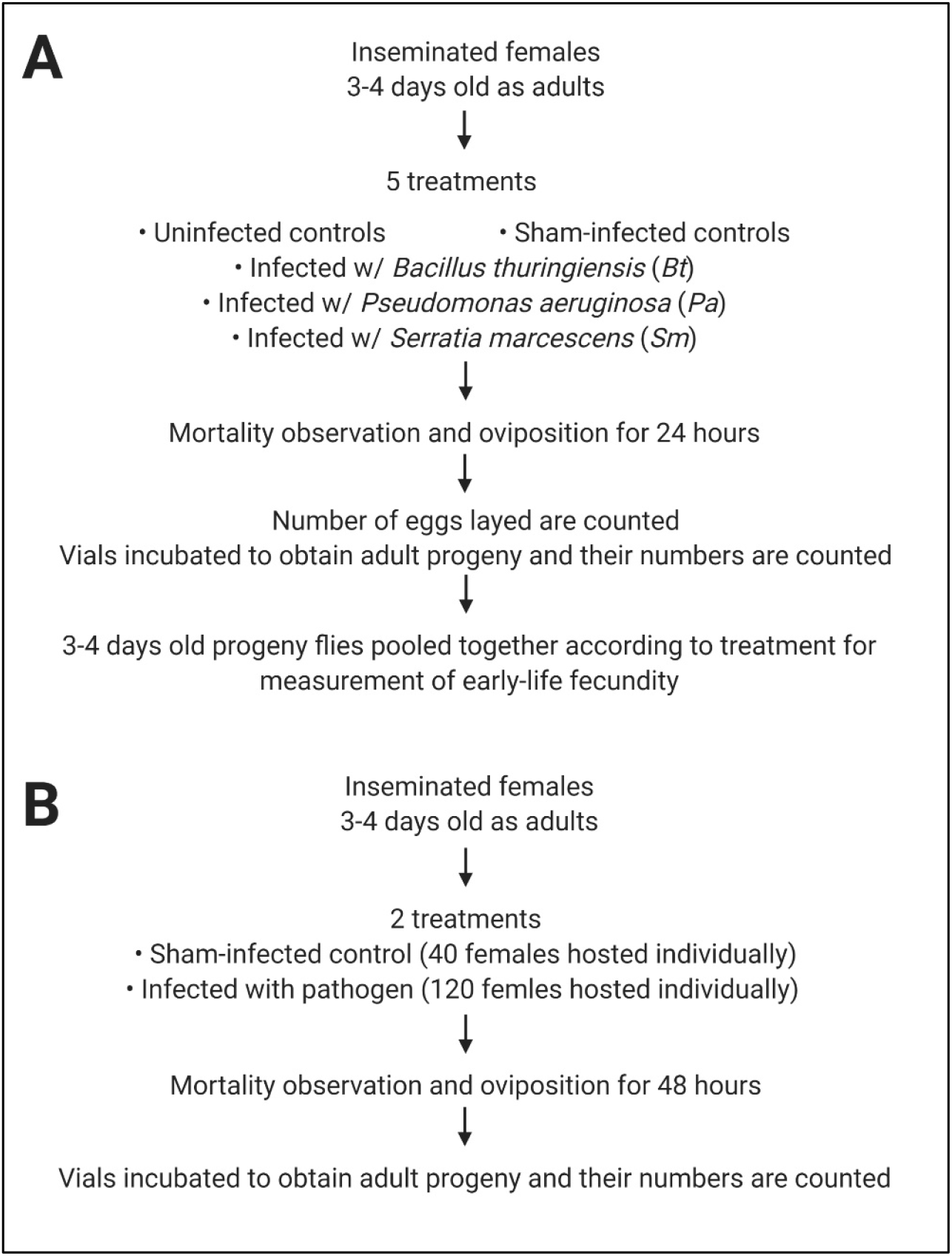
Experimental design for testing the effect of (A) pathogen identity, and (B) infection outcome and individual variability, on post-infection reproductive output of *Drosophila melanogaster* females.

All three pathogens used imposed significant mortality upon the infected females compared to the uninfected controls (figure 2.A). All females infected with *Sm* (hazard ratio, 95% confidence interval: 31895.33, 4333.172-234773.08) and *Pa* (HR, 95% CI: 936.66, 130.79-6707.86) died because of infection within 12 and 24 hours of infection, respectively, while about half of all females infected with *Bt* (HR, 95% CI: 156.17, 21.80-1118.65) died of infection within the observation period. Females that were sham-infected (HR, 95% CI: 7.98, 0.99-63.78) did not show significant difference in mortality compared to uninfected controls.

**Figure 2.**
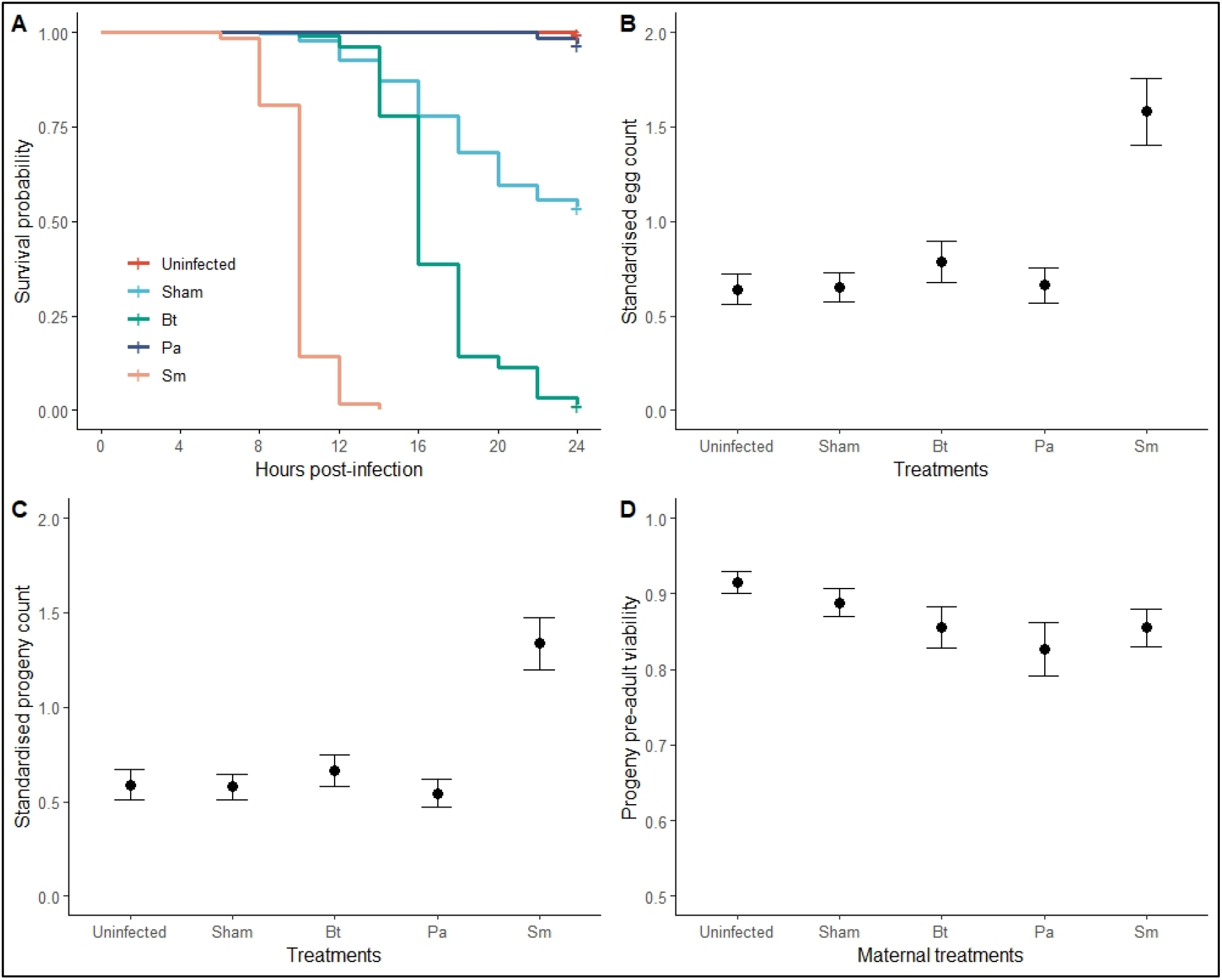
Effect of infection with different pathogens on (A) survival, (B) number of eggs produced, (C) number of progeny produced, and (D) pre-adult viability of progeny, of *Drosophila melanogaster* females.

Since the lifespan of females in each treatment was different from one another, instead of directly comparing the absolute number of eggs laid (or progeny produced), we divided the total number of eggs (or progeny) in each vial with the summation of the hours survived (survival time right-censored at 24 hours post-infection for surviving females) by the females in that vial (see MATERIALS AND METHODS for more details). We call this the “standardised reproductive output” and use this value as the subject of analysis. This value is essentially the measure of the number of eggs (or progeny) per female per hour, when the females are either infected with pathogens, or sham-infected, or left uninfected.

Infection treatment had a significant effect on standardised reproductive output, in terms of both eggs laid (F_4,147_: 58.778, p < 2.2 e-16; figure 2.B) and progeny produced (F_4,147_: 61.338, p < 2.2 e-16; figure 2.C). Post-hoc pairwise comparison using Tukey’s HSD indicated that *Sm*-infected females (least-square mean, 95% CI: 1.580, 1.408-1.752) laid a significantly greater number of eggs per female per hour compared to uninfected (LS mean, 95% CI: 0.641, 0.469-0.813), sham-infected (LS mean, 95% CI: 0.653, 0.481-0.825), *Bt*-infected (LS mean, 95% CI: 0.783, 0.611-0.955), and *Pa*-infected (LS mean, 95% CI: 0.662, 0.490-0.834) females; the other four experimental treatments did not differ from one another significantly in terms of number of eggs laid (table S2.A). Similarly, *Sm*-infected females (LS mean, 95% CI: 1.337, 1.207-1.467) produced a significantly greater number of progeny per female per hour compared to uninfected (LS mean, 95% CI: 0.589, 0.458-0.719), sham-infected (LS mean, 95% CI: 0.578, 0.448-0.708), *Bt*-infected (LS mean, 95% CI: 0.663, 0.533-0.794), and *Pa*-infected (LS mean, 95% CI: 0.545, 0.414-0.675) females; the other four experimental treatments did not differ from one another significantly in terms of number of progeny produced (table S2.B).

Infection treatment had a significant effect on progeny pre-adult viability (F_4,150_: 7.985, p = 7.304 e-06; figure 2.D). Post-hoc pairwise comparison using Tukey’s HSD indicated that progeny of *Bt*-infected (LS mean, 95% CI: 0.856, 0.830-0.881), *Pa*-infected (LS mean, 95% CI: 0.826, 0.800-0.851), and *Sm*-infected (LS mean, 95% CI: 0.855, 0.829-0.880) females had significantly less pre-adult viability compared to progeny of uninfected females (LS mean, 95% CI: 0.915, 0.889-0.940). There was no difference in viability between progeny of sham-infected (LS mean, 95% CI: 0.888, 0.863-0.914) and uninfected females. Progeny of *Pa*-infected females also had less viable compared to progeny of sham-infected females (table S2.C).

To have an estimate of the effect of infecting females with different pathogenic bacteria on the fitness of their progeny, we measured the early-life fecundity of the progeny, beginning at day 4-5 of adulthood till day 10-11 of adulthood. We pooled all progenies produced by all 80 females in each treatment (8 females × 10 vials) and randomly samples 60 males and 60 females, housing them in groups of 6 males and 6 females, setting up 10 vials per *maternal* treatment. We counted the number of progenies produced by these flies over the next six days, counting the progenies per day separately, and using that as the subject of analysis. The day of count (age of the flies) had a significant effect on early-life fecundity of progeny flies (F_1,897_: 488.713, p < 2.2 e-16; figure S1). Progeny early life fecundity was also significantly affected by *maternal* treatment (F_4,897_: 31.427, p < 2.2 e-16; figure S1). Post-hoc pairwise comparison using Tukey’s HSD indicated that progeny of *Sm*-infected females (LS mean, 95% CI: 9.56, 8.52-10.06) had significantly lesser fecundity compared to progeny of uninfected (LS mean, 95% CI: 12.08, 11.04-13.1), sham-infected (LS mean, 95% CI: 12.06, 11.02-13.1), *Bt*-infected (LS mean, 95% CI: 11.59, 10.55-12.6), and *Pa*-infected (LS mean, 95% CI: 11.79, 10.75-12.8) females; the other four *maternal* treatments did not differ from one another significantly in terms of progeny fecundity (table S2.D).

In the above experiment, all females infected with *Sm* and *Pa* died of infection, while only half of *Bt*-infected females died (figure 2.A). In the follow-up experiment (figure 1.B), we tested if the outcome of infection (survival vs. death), and the time of death for individual females, had any effect on reproductive fitness of the females. We housed infected and sham-infected females individually in food vials after infection, and monitored their mortality every 2 hours for 48 hours post-infection. The experiment was independently replicated thrice for each bacterium used: *Bt, Pa*, and *Sm*. We counted the number of progeny produced by individual females in the span of 48 hours (or till the time the female died) as a measure of reproductive output. To account for differences in lifespan (number of hours survived by an infected female; survival time right-censored at 48 hours post-infection for females that didn’t die within that time), we divided the number of progeny produced by a female by the hours survived, and used this “standardised reproductive output” as subject of analysis. Since in the first experiment, between treatment differences in standardised reproductive output did not change based on whether we focused on the number of eggs or the number of progeny, in this experiment we only counted the number of progeny produced.

Similar to the first experiment, only about half of *Bt*-infected females, while all of *Sm*- and *Pa*-infected females, died due to infection (figure 3.A, 3.D, 3.G). For *Bt*-infected females, infection outcome did not have a significant effect on mean standardised reproductive output of the females (F_2,475_: 1.4701, p = 0.2309); the infected-dead females (LS mean, 95% CI: 0.926, 0.683-1.17), the infected-alive females (LS mean, 95% CI: 0.895, 0.654-1.14), and the sham-infected females (LS mean, 95% CI: 0.817, 0.577-1.06) had comparable mean standardised reproductive output (figure 3.B). Infection outcome significantly affected the variance in standardised reproductive output (Levene’s test, F_2,475_: 20.808, p = 2.174 e-09), with infected-dead females exhibiting greater variance compared to both infected-alive and sham-infected females (figure 3.B). Within infected-dead female, time-of-death had a significant effect on standardised reproductive output (F_1,210_: 6.3233, p = 0.01267), with reproductive output having a mild negative correlation with time-to-death (coefficient, 95% CI: -0.01733, -0.03109 – -0.00355; η^2^, 90% CI: 0.03, 0.00-0.08; figure 3.C).

**Figure 3.**
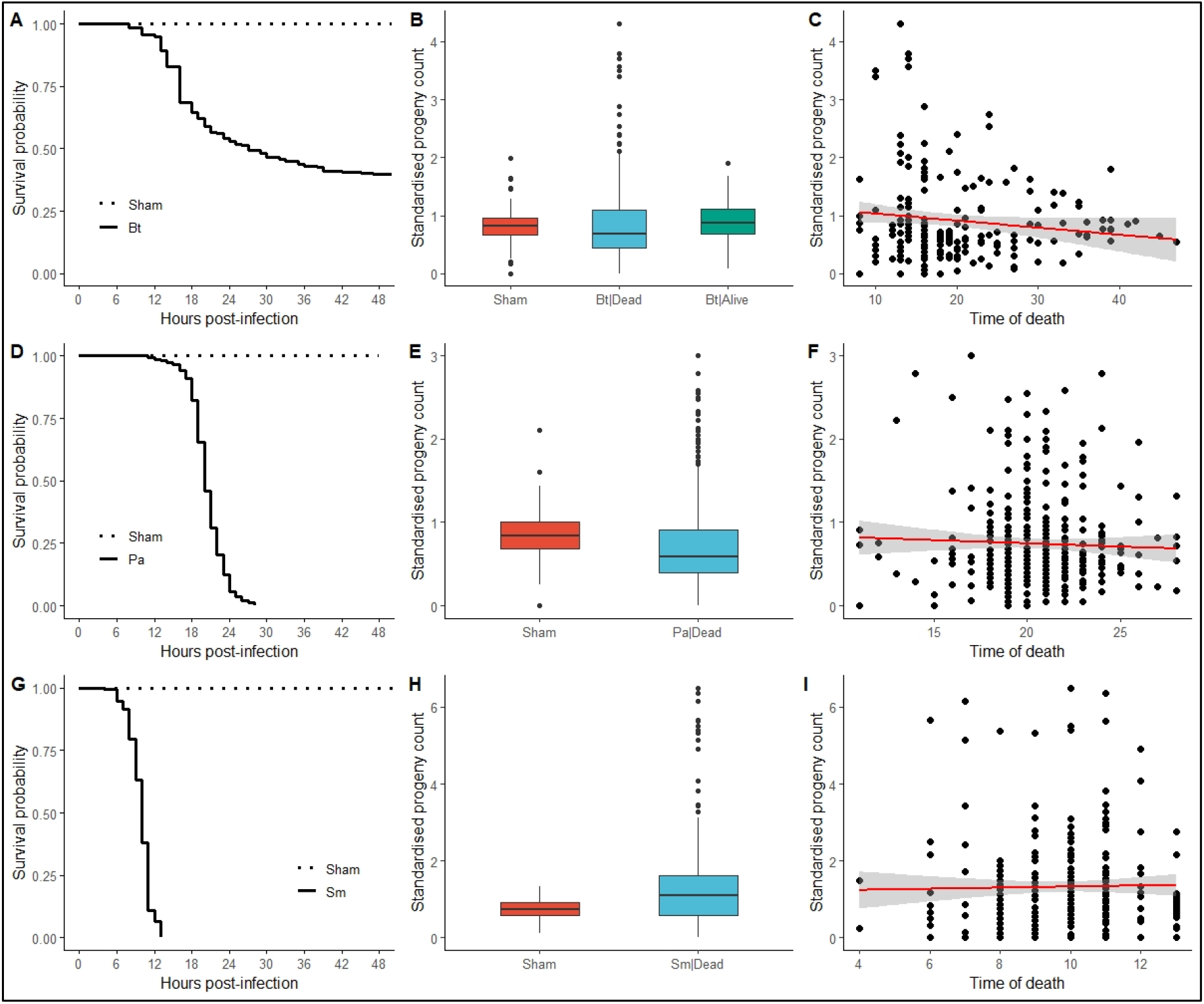
Effect of infection outcome on post-infection reproductive output of *Drosophila melanogaster* females, infected with *Bacillus thuringiensis* (A: survival, B: effect of infection outcome on progeny count, C: effect of time of death on progeny count), *Pseudomonas aeruginosa* (D: survival, E: effect of infection outcome on progeny count, F: effect of time of death on progeny count), and *Serratia marcescens* (G: survival, H: effect of infection outcome on progeny count, I: effect of time of death on progeny count).

For *Pa*-infected females (all of which died following infection; figure 3.D), infection outcome had a significant effect on standardised reproductive output (F_1,474_: 4.3739, p = 0.03703), with infected-dead females (LS mean, 95% CI: 0.743, 0.618-0.867) producing less number of progeny compared to sham-infected females (LS mean, 95% CI: 0.851, 0.721-0.981; figure 3.E). Infection outcome significantly affected the variance in standardised reproductive output (Levene’s test, F_2,474_: 19.795, p = 1.075 e-05), with infected-dead females exhibiting greater variance compared to sham-infected females (figure 3.E). Time-of-death did not have a significant effect on standardised reproductive output (F_1,357_: 0.511, p = 0.4503; figure 3.F).

For *Sm*-infected females (all of which died following infection; figure 3.G), infection outcome had a significant effect on standardised reproductive output (F_1,408_: 25.5, p = 6.684 e-07), with infected-dead females (LS mean, 95% CI: 1.28, 0.747-1.81) producing greater number of progeny compared to sham-infected females (LS mean, 95% CI: 0.72, 0.207-1.23; figure 3.H). Infection outcome significantly affected the variance in standardised reproductive output (Levene’s test, F_2,408_: 40.875, p = 4.444 e-10), with infected-dead females exhibiting greater variance compared to sham-infected females (figure 3.H). Time-of-death did not have a significant effect on standardised reproductive output (F_1,290_: 0.1505, p = 0.6983; figure 3.I).

## DISCUSSION

Fecundity compensation (or, terminal investment) theory in its simplest form hypothesises that an infected host facing impending death would increase its immediate reproductive effort to compensate for the loss of future opportunities to reproduce [1, 6, 7]. This hypothesis can be sub-structured into testable predictions, such as

a. hosts infected with a lethal pathogen would exhibit increased reproductive effort compared to hosts infected with a pathogen that does not kill all of the infected individuals;
b. in case of pathogens for which all hosts do not succumb to infection, hosts that die due to infection would increase their reproductive effort compared to hosts that survive the infection; and,
c. among hosts that succumb to infection, there will be a negative correlation between reproductive effort and time of death.

Tests of theoretical predictions of change in investment towards reproduction, in response to various intrinsic and extrinsic variables, hinge on accurate estimation of reproductive effort (proportion of total available resources that is invested towards reproduction), which is often difficult to measure [35]. Studies exploring infection induced changes in reproductive investment subvert this problem using uninfected controls. The uninfected controls represent an optimal reproductive output given a certain level of accessible resources and residual reproductive value. Resultantly in such studies a change in reproductive output in infected hosts compared to controls can be interpreted in light of the fecundity compensation/terminal investment theory [1, 6, 7]. In this study we use ‘*standardised* reproductive output’ (number of eggs, or progeny, normalised by the post-infection time-to-death of the females) as a proxy of reproductive effort. The lifespan of infected females in our experiments vary greatly depending upon the pathogen used for infection (figure 2.A), and thus a direct comparison of absolute egg or progeny count is not suitable.

Briefly, in this study we investigated how infection with three entomopathogenic bacteria, which differ from one another with respect to the level of mortality imposed on the host, affect the reproductive output of female *Drosophila melanogaster*. Additionally, we explored the effect of *maternal* infection on pre-adult viability and early-life fecundity of the progeny. We further investigated if infection outcome (death vs. survival), and the time of death, differentially affected the reproductive output of individual infected females. Our key findings are as follows:

a. The effect of infection on mean reproductive output is pathogen dependent (figures 2.B-C). Females infected with *Serratia marcescens* (hereafter *Sm*) produce a greater number of eggs (and progeny) compared to uninfected control females, after accounting for differences in post-infection lifespan. Females infected with *Bacillus thuringiensis* (hereafter *Bt*) or *Pseudomonas aeruginosa* (hereafter *Pa*) have reproductive output similar to controls.
b. The effect of *maternal* infection on progeny life-history is different for each trait measured. Progeny pre-adult viability was reduced by infection with all three pathogens, with the greatest reduction seen in progeny of *Pa*-infected females (figure 2.D). On the other hand, progeny early-life fecundity was compromised only in case of progeny of *Sm*-infected females; progeny of *Bt*- and *Pa*-infected females had fecundity comparable to progeny of uninfected females (figure S1).
c. Females that succumb to infection exhibit greater variability in reproductive output, compared to control females and females that survive the infection, irrespective of the pathogen used for infection (figures 3.B, 3.E, and 3.H). This variability in reproductive output is not explained by time of death in *Pa*- and *Sm*-infected female (figures 3.F and 3.I); for *Bt*-infected females there is a negative correlation between time of death and *standardised* reproductive output, but with a very low effect size (figure 3.C). Females that survive the infection have reproductive output comparable to controls in terms of both mean and variance (comparison possible for *Bt*-infected females only).

Forbes [18] classified host-pathogen systems based on whether acute infection had any negative effect on current reproduction (possibly due to somatic damage to the reproductive tissue or leeching of resources) and future reproductive potential (brought about by host death or permanent somatic damage) of the host. Increased reproductive output is the predicted outcome only if future reproductive potential is compromised, but without any negative effect on current reproduction [18]. An observed reduction in reproductive output during acute infection can thus be because of (a) pathogen leeching resources from the host or manipulating host physiology to reduce fecundity [12], (b) damage to reproductive tissue by the pathogen [24] or by the host immune defence itself [14], or (c) rerouting of resources meant for reproduction towards immune defence by the host [31]; although such reallocation of resources in either direction may not always translate into greater fitness benefits [19].

Amongst the three pathogens used in this study, infection with two (*Pa* and *Sm*), is absolutely lethal, while about half of females infected with *Bt* survive acute infection (figure 2.A). Therefore, *Sm*- and *Pa*-infected females have zero future reproductive potential, while *Bt*-infected females can continue to reproduce post-recovery, assuming that there is no lingering somatic damage. *Drosophila melanogaster* flies never clear out infecting pathogens from their system completely [36], and a chronic, low level of pathogens continue to persist with in the fly body, which requires some investment into immune function on part of the host to keep in check [33]. It is therefore a possibility that *Bt*-infected females may never regain uninfected levels of reproduction even post-recovery, but given that *Bt*-infected females that survive the infection continue to reproduce at levels of control females even during acute infection period (figure 3.B), this is an unlikely possibility. *Bt*-infected females should therefore invest towards immune defence and not towards increasing immediate reproductive output, to maximise chances of survival and opportunity of future reproduction, as we see in the results from our experiment (figure 2.B and 2.C).

Based on the arguments outlined above it is expected that females would increase their reproductive effort after being infected with *Sm* and *Pa*, but we observe an increase in reproductive output only in case of *Sm*-infected females (figure 2.B and 2.C). The absence of any change in reproductive output of *Pa*-infected females may be driven by many possible reasons, including damage to reproductive tissue, and exploitation or manipulation of host by the pathogen; we rule out resource reallocation driven costs since we have argued above that when infection guarantees lethality, investment away from reproduction is counter-productive. Since we directly did not measure damage to reproductive tissue, we cannot choose with sanguinity between the different possibilities listed above based on the data at hand.

Progeny of infected females, independent of the infecting pathogen suffered from reduced pre-adult viability; progeny of *Pa*-infected females exhibited the greatest reduction (figure 2.D). Perrin et al [37] have proposed that when *maternal* infection compromises progeny viability, increasing progeny production is not a suitable strategy for an infected host; this may be another explanation for why *Pa*-infected females do not increase reproductive output despite of guaranteed lethality due to infection. Reduced viability of progeny of Pa-infected females have been reported in other previous studies (viz. [38]; but see [23]). Reduced progeny viability can lead to a progeny quantity vs. progeny quality trade-off [2], making investment into progeny quality, instead of increasing progeny number, a potential strategy for *Pa*-infected females.

Infection with *Pa* has been previously demonstrated to both increase [23] and suppress [25] reproduction in females. A host’s response to the same pathogen can change because of the route of infection [22, 29], which can be a possible reason behind different observations in different studies: Hudson et al [23] infected flies via oral route, while in this study and in that of Linder and Promislow [25] flies were infected via septic injury to the thorax. A systemic infection is more likely to reach the reproductive tissues, via the haemolymph, than an oral infection, which first has to colonise the gut and breach the gut lining to enter into circulation.

Infected females that died of infection, irrespective of the pathogen used for infection, exhibited greater inter-individual variability in reproductive output compared to control (sham-infected) females and females that survived the infection (figures 3.B, 3E, and 3.H). The observed difference in the mean reproductive outputs of the infected and control females remained consistent across both experiments, suggesting that pathogen identity is a reliable predictor of post-infection reproductive output at the population level. Females that survived the infection with *Bt* had reproductive output similar to that of controls (figure 3.B), in terms of both population mean and inter-individual variability, suggesting that infected females may be able to judge their own prognosis and invest into reproduction accordingly. Since no females infected with either *Pa* or *Sm* survived the infection, we cannot conclude if this observation is generalizable for all other pathogens for which mortality is less than hundred percent. What seems puzzling therefore is why don’t females that succumb to infection, irrespective of pathogen identity, increase their reproductive output?

Based on the earlier discussion, individual females that succumb to infection are expected to increase their reproductive output. This increase should happen irrespective of pathogen identity, driven only by the risk of mortality, except in a case where infection compromises current reproductive capacity. Contrary to this expectation, we see that the reproductive output of infected-dead females for each pathogen ranges from zero to extremely high values; with some females reproducing less compared to the controls and other reproducing far more in excess (figures 3.B, 3E, and 3.H). Therefore, in canonical sense, we see some females exhibiting ‘cost of immunity’ while other females exhibiting ‘fecundity compensation’ when infected with the very same pathogen. This variation in reproductive output seems to be independent of both pathogen identity and time of death.

The observed inter-individual variability in reproductive output of females that die of infection may purely be stochastic, without any consequence in terms of evolutionary outcomes [39]. Alternatively, the heterogeneity may reflect variation in individual female quality [40] and physiological state [41]. The physiological state of an individual is a potent predictor of its residual reproductive value, and all else being equal, can therefore influence infection-induced changes in reproductive effort [42]. A third possibility is that the heterogeneity is a consequence of host variation, genetic or otherwise, in response to infection, in terms of both resistance and tolerance [15, 27, 32, 43]. Further empirical exploration is necessary to disentangle these potential causes of inter-individual variation.

To summarize, we find that lethal infections do not always induce an increased investment towards immediate reproduction in female *Drosophila melanogaster*; females infected with only one out of two pathogens that imposed hundred percent mortality increased their reproductive effort. Furthermore, females dying of infection do not have greater reproductive effort compared to females that survive the infection, and reproductive effort had a negative correlation with time of death in case only one out of three pathogens used in this study. These findings suggest that the mechanistic interaction between a host and a pathogen has a greater influence on host reproductive effort, compared to infection status, infection outcome, and mortality risk on the host by the pathogen. Additionally, our results suggest that pathogen identity is a reliable predictor of bacterial infection induced change in reproductive effort of the females at the level of population means, but pathogen identity does not predict reproductive output of individual females. Females infected with all three pathogens used in this study have overlapping range of reproductive output. Furthermore, maternal infection can affect progeny life-history traits, but the effect is specific to individual traits. In conclusion, dichotomy of ‘cost of immunity’ versus ‘fecundity compensation’ is too narrow in scope to account for all nuances involved in post-infection change in reproductive effort.

## MATERIALS AND METHODS

### Host population and general handling

Flies from BRB2 population - a large, lab adapted, out-bread population of Drosophila melanogaster - was used for the experiments reported in this paper. The <u>B</u>lue <u>R</u>idge <u>B</u>aseline (BRB) populations were originally established by hybridizing 19 wild-caught iso-female lines [44], and has been maintained since then as an outbred population on a 14-day discrete generation cycle with census size of about 2800 adults each generation. Every generation, eggs are collected from population cages (plexiglass cages: 25 cm length × 20 cm width × 15 cm height) and dispensed into vials (25 mm diameter × 90 mm height) with 8 ml banana-jaggery-yeast food medium, at a density of 70 eggs per vial. 40 such vials are set up; the day of egg collection is demarcated as day 1. The vials are incubated at 25 °C, 50-60% RH, 12:12 hour LD cycle; under these conditions the egg-to-adult development time for these flies is about 9-10 days. On day 12 post egg collection all adults are transferred to population cage, and provided with fresh food plates (banana-jaggery-yeast food medium in a 90 mm Petri plate) supplemented with *ad libitum* live yeast paste. On day 14, the cage is provided with fresh food plate, and 18 hours later eggs are collected from this plate to begin the next generation.

### Pathogen handling and infection protocol

Three bacterial pathogens were used in this study for infecting the flies: *Bacillus thuringiensis* (*Bt*; obtained from DSMZ, Germany, catalogue number: DSM2046), *Pseudomonas aeruginosa* (*Pa*; obtained from MTCC, India), and *Serratia marcescens* (*Sm*; [29]). All three pathogens are maintained in the lab as glycerol stocks, and are cultured in Luria Bertani broth (Himedia, M1245); cultures are incubated at 30 °C for *Bt*, and 37 °C for *Pa* and *Sm*. Overnight culture of bacteria grown from glycerol stocks was diluted (1:100) in fresh LB medium and incubated till confluency (optical density OD_600_ = 1.0-1.2). The bacterial cells were pelleted down by centrifugation and re-suspended in sterile 10 mM MgSO_4_ buffer at OD_600_ = 1.0. Flies were infected by pricking them at the dorsolateral side of the thorax with a fine needle (Minutien pin, 0.1 mm, Fine Science Tools, CA, item no. 26001-10) dipped in bacterial suspension under light CO_2_ anesthesia. Flies for sham-infections were similarly treated, but pricked with needle dipped in sterile 10 mM MgSO_4_ buffer. Uninfected control flies were only subjected to CO_2_ anesthesia.

### Generation of experimental flies

Eggs were collected from BRB2 population cages and distributed into food vials with 8 ml of standard food medium at a density of 70 eggs per vial. These vials were incubated as per general maintenance. Twelve days post egg-laying flies were flipped into fresh food vials and hosted for two more days before experimentation. This ensured all focal females were 4-5 day old, sexually mature and inseminated, at the time of infections. Flies were again flipped into fresh food vials 6 hours before being subjected to experimental treatments (as described below).

### Experimental design

#### Experiment 1

Focal females were randomly distributed into five treatments: (a) infected with *Bacillus thuringiensis* (*Bt*), (b) infected with *Pseudomonas aeruginosa* (*Pa*), (c) infected with *Serratia marcescens* (*Sm*), (d) sham-infected controls, and (e) uninfected controls. The entire experiment was independently replicated thrice. Flies were placed in fresh food vials after being subjected to respective treatments. For each treatment 10 vials were set up, each with 8 females for oviposition; each vial was used as a unit of replication. The vials were monitored every 2 hours to record any mortality, for 24 hours post-infection, divided into two consecutive 12-hour windows. Flies alive at the end of first 12-hour window were flipped into fresh food vials (one-to-one mapping of vial identity), and flies alive at the end of 24 hours were discarded (censored). The number of eggs in each vial was counted at the end of respective 12-hour windows. The vials were then incubated under standard maintenance conditions for the eggs to develop into adults, and 12 days after the oviposition period, all adult progeny were counted under light CO_2_ anesthesia and transferred to fresh food vials.

*Standardized reproductive output* of females in each vial was calculated as,

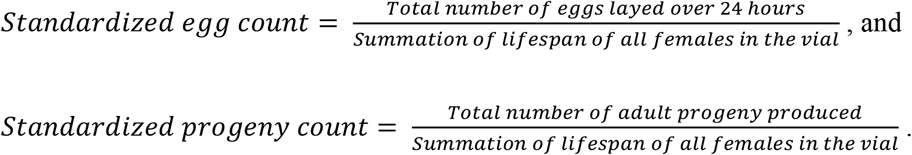

Standardization was carried out to account for the differences in post-infection survival time of females in various treatments (see RESULTS for more details). *Progeny pre-adult viability* for each vial was calculated as,

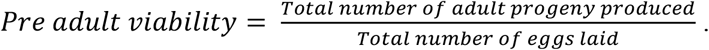

Following progeny counts, 4-5-day old adult progeny were pooled together according to treatments and distributed to fresh food vials with 5 females and 5 males in each vial; 10 such vials were set up per *maternal* treatment per replicate. These flies were allowed to oviposit for six consecutive days (by flipping them into fresh food vials every day) to obtain an estimate of *offspring early-life fecundity*. These vials were incubated at standard maintenance conditions, and the number of progeny in these vials were counted 12 days post-oviposition.

#### Experiment 2

Focal females were randomly distributed into two treatments: (a) infected with bacteria, and (b) sham-infected controls. For infected treatment, 120 females were individually hosted in vials for oviposition, while for sham-infected controls 40 females were hosted individually. The experiment was replicated thrice with each pathogen. (Due to a handling accident, one replicate with Sm had sample size of 60 and 30 females for infected and sham-infected treatments, respectively.) The vials were monitored every 2 hours for any mortality, for 48 hours post-infection, after which the alive flies were discarded. The vials were then incubated under standard maintenance conditions for the eggs to develop into adults, and 12 days later the number of adult progeny was counted for each individual female.

*Standardized reproductive output* for each individual female was calculated as,

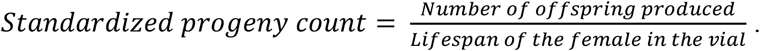

### Statistical analysis

All analyses were carried out using R statistical software (version 4.1.0 [45]), using various functions from the *survival* [46], *coxme* [47], *lmerTest* [48], *emmeans* [49], and *car* [50] packages. Graphs were created using the *ggplot2* [51] and *survminer* [52].

#### Experiment 1

Survival data was analyzed using mixed-effects Cox proportional hazards model, with ‘Treatment’ as a fixed factor and ‘Replicate’ as a random factor. Reproductive output, progeny viability, and progeny early-life fecundity was modeled using linear models (as described below) and subjected to significance testing using type III analysis of variance (ANOVA). Post-hoc pairwise comparisons, wherever necessary, was carried using Tukey’s HSD method.

Standardized egg count ∼ Treatment + (1|Replicate)

Standardized progeny count ∼ Treatment + (1|Replicate)

Progeny pre-adult viability ∼ Maternal treatment + (1|Replicate)

Progeny early-life fecundity ∼ Day + Maternal treatment + (1|Replicate)

Significance tests for random effects are tabulated in table S1.

#### Experiment 2

Reproductive output data was modeled using a linear model (as described below) and subjected to significance testing using type III ANOVA. Post-hoc pairwise comparisons, wherever necessary, was carried using Tukey’s HSD method.

Standardized progeny count ∼ Category + (1|Replicate)

‘Category’ denoted the combination of infection status and infection outcome, and consists of three levels: sham-infected females, infected-alive females, and infected-dead females. Effect of time on death on reproductive output of infected-dead females was similarly analyzed with type III ANOVA using the following linear model:

Standardized progeny count ∼ Time of death + (1|Replicate)

Significance tests for random effects are tabulated in table S1. Comparison of variances across ‘category’ was carried out using Levene’s test after pooling data from all three replicates for each pathogen.

## Acknowledgements

We thank Paresh Nath Das for logistical support during execution of the experiments reported in this manuscript. We also thank Aparajita Singh and Rochishnu Dutta for their constructive comments on earlier drafts of the manuscript. We thank Dr. E Sucena and Tania Paulo (Instituto Gulbenkian Ciencia, Portugal) for providing us with the *Serratia marcescens* isolate used in experiments reported in this manuscript.

## SUPPLEMENTARY FILES

**Figure S1.**
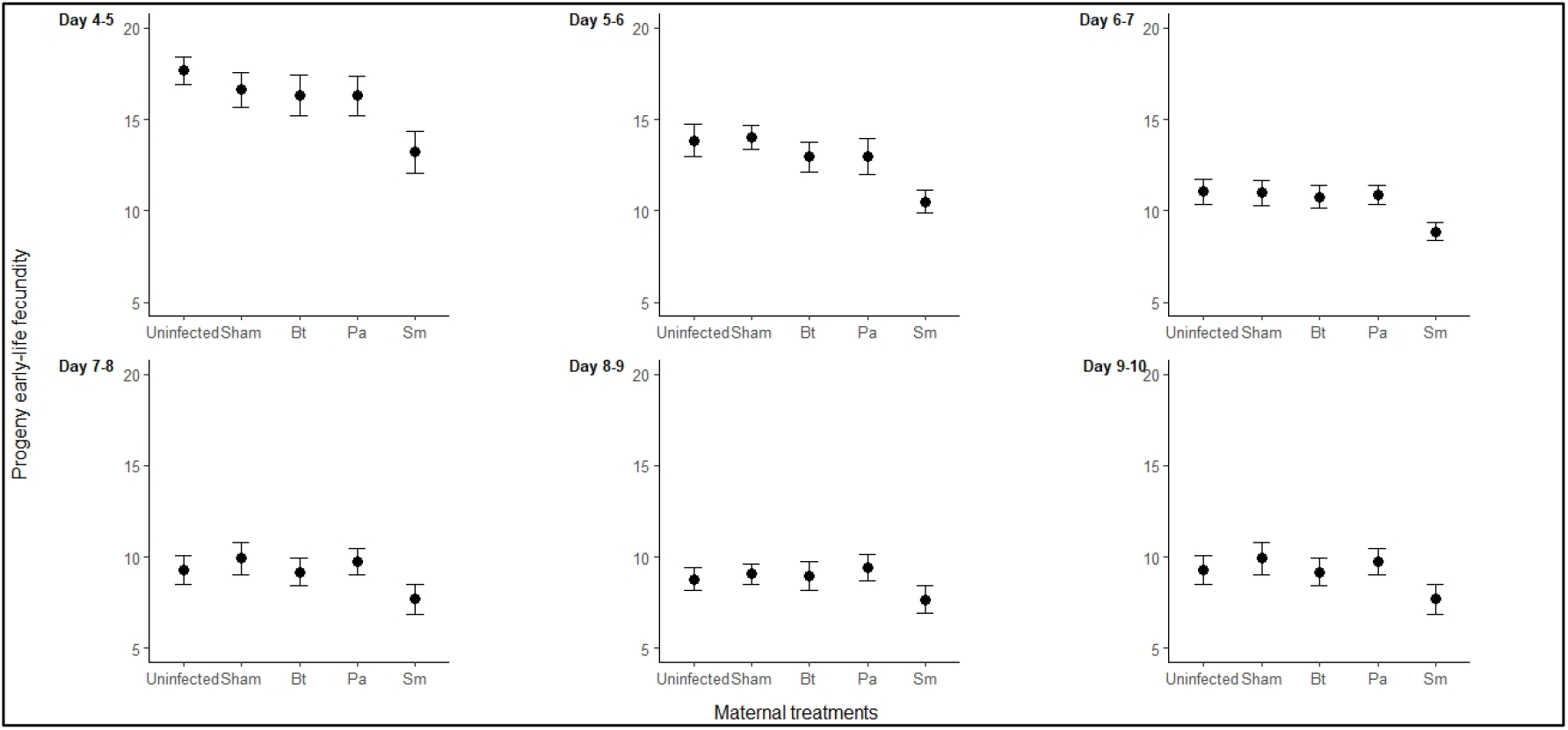
Effect of *maternal* infection treatment on progeny early-life fecundity; fecundity measured on each day shown separately.

**Table S1**. Significance tests for random factors included in various type-III ANOVA reported in the ‘Results’ section. See ‘Materials and Methods’ for full details on statistical analysis.

**Table S2**. Post-hoc pairwise comparisons using Tukey’s HSD for significant effects reported for fixed factors in various type-III ANOVA reported in the ‘Results’ section. See ‘Materials and Methods’ for full details on statistical analysis.

