## Supplemental Table 1 for "Pathogen dependence and inter-individual variability of post-infection reproductive fitness in *Drosophila melanogaster*"

**Table S01.** Significance tests for random effects included in type III ANOVA reported in text.

| Factors | npar | logLik | AIC | LRT | Df | Pr (>Chisq) |
|---|---|---|---|---|---|---|
| **(A) Effect of infection treatment on standardized egg count (experiment 1)** | | | | | | |
| <none> | 7 | -29.068 | 72.137 | | | |
| (1\|Replicate) | 6 | -32.425 | 76.850 | 6.7136 | 1 | 0.009568 |
| **(B) Effect of infection treatment on standardized progeny count (experiment 1)** | | | | | | |
| <none> | 7 | 2.6005 | 8.799 | | | |
| (1\|Replicate) | 6 | 0.1193 | 11.761 | 4.9624 | 1 | 0.0259 |
| **(C) Effect of *maternal* infection treatment on progeny pre-adult viability (experiment 1)** | | | | | | |
| <none> | 7 | 194.06 | -374.11 | | | |
| (1\|Replicate) | 6 | 194.06 | -376.11 | 0 | 1 | 1 |
| **(D) Effect of *maternal* infection treatment on progeny early-life fecundity (experiment 1)** | | | | | | |
| <none> | 8 | -2114.8 | 4245.5 | | | |
| (1\|Block) | 7 | -2132.0 | 4278.0 | 34.465 | 1 | 4.34e-09 |
| **(E) Effect of infection outcome on standardized progeny count; *B. thuringiensis* (experiment 2)** | | | | | | |
| <none> | 5 | -401.31 | 812.62 | | | |
| (1\|Replicate) | 4 | -409.76 | 827.53 | 16.904 | 1 | 3.931e-05 |
| **(F) Effect of time of death on standardized progeny count; *B. thuringiensis* (experiment 2)** | | | | | | |
| <none> | 4 | -247.28 | 502.57 | | | |
| (1\|Replicate) | 3 | -250.08 | 506.16 | 5.5972 | 1 | 0.01799 |
| **(G) Effect of infection outcome on standardized progeny count; *P. aeruginosa* (experiment 2)** | | | | | | |
| <none> | 4 | -338.55 | 685.09 | | | |
| (1\|Replicate) | 3 | -339.93 | 685.87 | 2.7761 | 1 | 0.09568 |
| **(H) Effect of time of death on standardized progeny count; *P. aeruginosa* (experiment 2)** | | | | | | |
| <none> | 4 | -290.01 | 588.03 | | | |
| (1\|Replicate) | 3 | -291.08 | 588.17 | 2.1433 | 1 | 0.1432 |
| **(I) Effect of infection outcome on standardized progeny count; *S. marcescens* (experiment 2)** | | | | | | |
| <none> | 4 | -578.99 | 1166.0 | | | |
| (1\|Replicate) | 3 | -590.23 | 1186.5 | 22.467 | 1 | 2.138e-06 |
| **(J) Effect of time of death on standardized progeny count; *S. marcescens* (experiment 2)** | | | | | | |
| <none> | 4 | -466.83 | 941.67 | | | |
| (1\|Replicate) | 3 | -476.45 | 958.91 | 19.238 | 1 | 1.154e-05 |
