## Supplemental Table 2 for "Pathogen dependence and inter-individual variability of post-infection reproductive fitness in *Drosophila melanogaster*"

**Table S02.** Post-hoc analysis using Tukey's HSD for pairwise comparisons (experiment 1).

| Pairwise comparison | Estimate | SE | DF | t-ratio | p-value |
|---|---|---|---|---|---|
| **(A) Effect of infection treatment on standardized egg count** | | | | | |
| Uninfected - Bt | -0.14222 | 0.0757 | 151 | -1.880 | 0.3327 |
| Uninfected - Pa | -0.02071 | 0.0757 | 151 | -0.274 | 0.9988 |
| Uninfected - Sham | -0.01165 | 0.0757 | 151 | -0.154 | 0.9999 |
| **Uninfected - Sm** | **-0.93912** | **0.0757** | **151** | **-12.412** | **<.0001** |
| Bt - Pa | 0.12151 | 0.0757 | 151 | 1.606 | 0.4960 |
| Bt - Sham | 0.13057 | 0.0757 | 151 | 1.726 | 0.4214 |
| **Bt - Sm** | **-0.79690** | **0.0757** | **151** | **-10.532** | **<.0001** |
| Pa - Sham | 0.00906 | 0.0757 | 151 | 0.120 | 1.0000 |
| **Pa - Sm** | **-0.91841** | **0.0757** | **151** | **-12.138** | **<.0001** |
| **Sham - Sm** | **-0.92747** | **0.0757** | **151** | **-12.258** | **<.0001** |
| **(B) Effect of infection treatment on standardized progeny count** | | | | | |
| Uninfected - Bt | -0.0749 | 0.0614 | 151 | -1.220 | 0.7399 |
| Uninfected - Pa | 0.0439 | 0.0614 | 151 | 0.715 | 0.9528 |
| Uninfected - Sham | 0.0108 | 0.0614 | 151 | 0.175 | 0.9998 |
| **Uninfected - Sm** | **-0.7483** | **0.0614** | **151** | **-12.193** | **<.0001** |
| Bt - Pa | 0.1187 | 0.0614 | 151 | 1.935 | 0.3035 |
| Bt - Sham | 0.0856 | 0.0614 | 151 | 1.395 | 0.6318 |
| **Bt - Sm** | **-0.6734** | **0.0614** | **151** | **-10.973** | **<.0001** |
| Pa - Sham | -0.0331 | 0.0614 | 151 | -0.540 | 0.9830 |
| **Pa - Sm** | **-0.7922** | **0.0614** | **151** | **-12.908** | **<.0001** |
| **Sham - Sm** | **-0.7591** | **0.0614** | **151** | **-12.368** | **<.0001** |
| **(C) Effect of *maternal* infection treatment on progeny pre-adult viability** | | | | | |
| **Uninfected - Bt** | **0.059137** | **0.0174** | **151** | **3.403** | **0.0075** |
| **Uninfected - Pa** | **0.088922** | **0.0174** | **151** | **5.117** | **<.0001** |
| Uninfected - Sham | 0.026788 | 0.0174 | 151 | 1.542 | 0.5372 |
| **Uninfected - Sm** | **0.059889** | **0.0174** | **151** | **3.446** | **0.0065** |
| Bt - Pa | 0.029786 | 0.0174 | 151 | 1.714 | 0.4285 |
| Bt - Sham | -0.032349 | 0.0174 | 151 | -1.862 | 0.3426 |
| Bt - Sm | 0.000752 | 0.0174 | 151 | 0.043 | 1.0000 |
| **Pa - Sham** | **-0.062134** | **0.0174** | **151** | **-3.576** | **0.0042** |
| Pa - Sm | -0.029033 | 0.0174 | 151 | -1.671 | 0.4551 |
| Sham - Sm | 0.033101 | 0.0174 | 151 | 1.905 | 0.3192 |

| (D) Effect of *maternal* infection treatment on progeny early-life fecundity | | | | | |
|---|---|---|---|---|---|
| Uninfected - Bt | 0.4849 | 0.267 | 902 | 1.817 | 0.3643 |
| Uninfected - Pa | 0.2900 | 0.267 | 902 | 1.086 | 0.8136 |
| **Uninfected - Sm** | **2.5128** | **0.267** | **902** | **9.415** | **<.0001** |
| Uninfected - Sham | 0.0146 | 0.267 | 902 | 0.055 | 1.0000 |
| Bt - Pa | -0.1950 | 0.267 | 902 | -0.730 | 0.9494 |
| **Bt - Sm** | **2.0278** | **0.267** | **902** | **7.598** | **<.0001** |
| Bt - Sham | -0.4703 | 0.267 | 902 | -1.762 | 0.3965 |
| **Pa - Sm** | **2.2228** | **0.267** | **902** | **8.328** | **<.0001** |
| Pa - Sham | -0.2754 | 0.267 | 902 | -1.032 | 0.8407 |
| **Sm - Sham** | **-2.4982** | **0.267** | **902** | **-9.360** | **<.0001** |
